# Peptidoglycan from *Bacillus anthracis Inhibits* Human Macrophage Efferocytosis in Part by Reducing Cell Surface Expression of MERTK and TIM-3

**DOI:** 10.1101/2023.03.30.535001

**Authors:** Joshua S Mytych, Zijian Pan, Charmaine Lopez-Davis, Nancy Redinger, Christina Lawrence, Jadith Ziegler, Narcis I. Popescu, Judith A. James, A. Darise Farris

## Abstract

*Bacillus anthracis* peptidoglycan (PGN) is a major component of the bacterial cell wall and a key pathogen-associated molecular pattern (PAMP) contributing to anthrax pathology, including organ dysfunction and coagulopathy. Increases in apoptotic lymphocytes are a late-stage feature of anthrax and sepsis, suggesting there is a defect in apoptotic clearance. Here, we tested the hypothesis that *B. anthracis* PGN inhibits the capacity of human monocyte-derived macrophages (MΦ) to efferocytose apoptotic cells. Exposure of CD163^+^CD206^+^ MΦ to PGN for 24h impaired efferocytosis in a manner dependent on human serum opsonins but independent of complement component C3. PGN treatment reduced cell surface expression of the pro-efferocytic signaling receptors MERTK, TYRO3, AXL, integrin αVβ5, CD36 and TIM-3, whereas TIM-1, αVβ3, CD300b, CD300f, STABILIN-1 and STABILIN-2 were unaffected. ADAM17 is a major membrane-bound protease implicated in mediating efferocytotic receptor cleavage. We found multiple ADAM17-mediated substrates increased in PGN-treated supernatant suggesting involvement of membrane-bound proteases. ADAM17 inhibitors TAPI-0 and Marimastat prevented TNF release, indicating effective protease inhibition, and modestly increased cell-surface levels of MerTK and TIM-3 but only partially restored efferocytic capacity by PGN-treated MΦ. We conclude that human serum factors are required for optimal recognition of PGN by human MΦ and that *B. anthracis* PGN inhibits efferocytosis in part by reducing cell surface expression of MERTK and TIM-3.

## INTRODUCTION

*Bacillus anthracis*, the causative agent of anthrax infection, is a large, spore-forming and toxin-producing gram-positive organism that typically infects herbivores. Humans can be infected through various routes including inhalation, ingestion or cutaneous exposure to spores, the most severe being inhalation which can lead to a rapid and highly fatal infection^1^. Latent spores migrate to and germinate in the mediastinal lymph nodes, followed by progression to fulminant infection characterized by high levels of circulating bacteria and sepsis-like features^2^. Bacterial sepsis is a leading cause of mortality worldwide and has limited treatment options^3^. Accumulation of uncleared apoptotic lymphocytes is a known hallmark of sepsis^4^, and is accompanied by increased levels of circulating nucleosomes^5^. Circulating nucleosomes can be released from uncleared apoptotic cells that progress to secondary necrosis^6^, promoting micro-and macro-vascular leakage^7^ and contributing to organ dysfunction^8^.

Macrophage (MΦ), either tissue-resident or those present in secondary lymphoid organs, are responsible for the clearance of apoptotic cells through a process termed efferocytosis^9-11^. Highly efferocytic MΦ tend to have an M2-like, or alternative activation program^12, 13^, expressing CD163 and CD206^14^ and acquire anti-inflammatory properties that actively dampen inflammation by secreting pro-resolving mediators^15^. Highly efferocytic MΦ can be modeled *in vitro* by polarization with IL-10, IL-4 and/or dexamethasone^16, 17^. Defects in efferocytosis amplify inflammation in sepsis by increasing the levels of circulating nucleosomes and other host damage-associated molecular patterns (DAMPs)^18^. Recognition of apoptotic cells is a complex process involving tethering receptors that on MΦ bind directly to phosphatidylserine on apoptotic cells or receptors that require soluble phosphatidylserine-binding bridge proteins^19^. Currently, there are at least twelve well-established receptors that signal for apoptotic cell engulfment: TYRO3, AXL, MERTK, integrins αVβ3 and αVβ5, CD300b, CD300f, STABILIN-1 and 2, TIM-1 and -3, and BAI-1^20^. Of these, TYRO3, AXL, and MERTK require bridging by GAS6, Protein S or GAL-3, and integrins αVβ3 and αVβ5 require bridging via MFGE8 or DEL-1. The remaining seven receptors directly bind to apoptotic cells^20, 21^. Efferocytic receptors are thought to be regulated at the cell surface by metalloproteases, and proteolysis of surface exposed TYRO3, AXL, MERTK, TIM-1, and TIM-3, has been reported in both mouse and human MΦ^22^. Loss of these receptors has been shown to correlate with disease phenotypes and may be useful as biomarkers in multiple diseases including lupus and Sjogren’s disease, where defects in efferocytosis are thought to contribute to disease progression^23, 24^. ADAM17 is a key sheddase responsible for cleaving more than 80 substrates, including TNF and MERTK, and has been implicated as a regulator of efferocytosis ^25, 26^.

We previously showed that *Bacillus anthracis* Edema Toxin, an adenylate cyclase that induces supraphysiologic levels of cAMP, impairs efferocytosis through inhibition of Rac1 activation and altered phosphorylation of proteins involved in actin reorganization^27^. In this study we focus on peptidoglycan (PGN), a major component of the cell wall in gram-positive bacteria and a significant factor in anthrax-mediated sepsis, that contributes to vascular and organ damage, and mortality in non-human primate models of anthrax sepsis^28, 29^. Response to PGN was initially thought to be at the cell surface and Toll-like receptor 2 (TLR2)-dependent^30^; however, recent work has challenged this view and showed that impurities from lipoteichoic acid (LTA) resulted in TLR-2 activation^31^. Further work elucidated that purified PGN preparations lacking LTA activate cytosolic Nucleotide-binding Oligomerization Domain (NOD) receptors^32-34^. Additional work using human monocytes, neutrophils and dendritic cells suggests that polymeric PGN activates NOD in human immune cells more potently in the presence of human serum opsonins^35-37^ however, this has yet to be shown for human MΦ. Herein, we demonstrate that recognition of *Bacillus anthracis* PGN by human MΦ is enhanced by human serum factors and that PGN inhibits human MΦ efferocytosis in part by reducing cell surface expression of MERTK and TIM-3.

## METHODS

### Differentiation of monocytes to M2-like MΦ

On Day 0 human mononuclear cells were isolated from fresh buffy coats using Lympholyte (Cedarlane labs, Burlington, Canada) according to the manufacturer’s protocol (see **Supplemental Figure 1** for timeline of assay). Following isolation, PBMC were counted and plated on 100mm petri dishes (Corning, NY, USA, 100mm × 20mm non-treated tissue culture dishes, Cat#: 430591) at 4×10^6^ cells/mL in 25mL per dish in Iscove’s Modified Dulbecco’s Medium with glutamate (IMDM) (Thermo Fisher Scientific, Waltham, MA, USA, Cat#:12440053) with no other additives. Cells were allowed to adhere for 1hr at 37°C in 5% CO_2_, followed by gentle removal of media and replacement with 20mL/dish of complete medium (IMDM, 10% hiFBS, 5mM PenStrep antibiotic) supplemented with 50ng/mL M-CSF (PeproTech, Rockyhill NJ, USA, Cat#: 300-25). On Days 2 and 5, half of the media was exchanged with fresh media containing 100ng/mL M-CSF to replenish spent M-CSF (final M-CSF concentration was 50ng/mL). On Day 6 MΦ were polarized to a tissue-like/alternative phenotype by adding 500nM dexamethasone and culturing for 24hrs (Sigma Aldrich, St. Louis, MO, USA, Cat#: D4902). All polarized MΦ expressed CD206 and CD163, as we previously published^27^.

**Figure 1:**
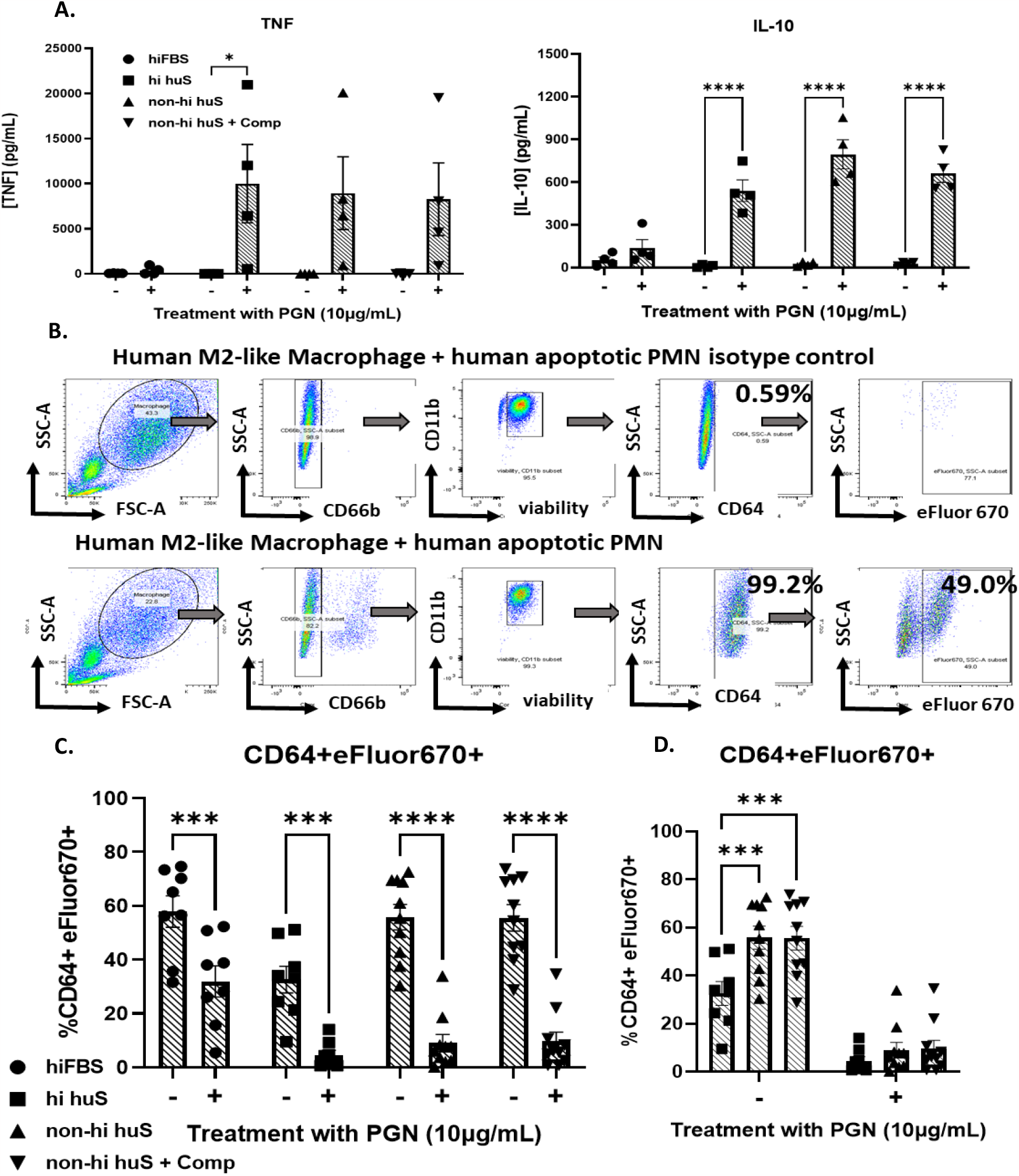
Bacillus anthracis peptidoglycan (PGN) inhibits efferocytosis in the presence of human serum. **A)** Secreted TNF and IL-10 (24hr-treated MΦ, in various serum conditions) from 4 independent donors. **B)** Gating strategy for flow cytometric evaluation of efferocytosis **C)** Percentage of CD64+ MΦ containing apoptotic eFluor670-labeled PMN from the indicated serum conditions in the presence or absence of 10ug/mL PGN from ≥5 independent donors analyzed by two-way ANOVA with Sidak’s multiple comparison. **D)** Alternative display of human serum condition data from Figure 1C to demonstrate hi huS has lower efferocytosis compared to the other human serum in untreated MΦ.

### PGN Treatment

On Day 7, 24hrs after polarization with dexamethasone, MΦ were re-plated to 12-well Nunc UpCell plates (Thermo Fisher, Cat#: 150200). Briefly, 3mL trypsin/EDTA solution (Sigma Aldrich, Louis, MO, USA, Cat#: T4049-100) was added to each petri dish and incubated at 37°C in 5% CO_2_ for 10 min to detach cells. After detachment, 10mL complete medium (containing 10%FBS) was added to neutralize trypsin activity. MΦ were pelleted at 1200 rpm for 5 min, counted and re-plated to the UpCell plates at 0.5×10^6^ cells/mL. After re-plating, MΦ were incubated for 16 or 24hrs with 10μg/mL highly purified *Bacillus anthracis* PGN^38^ (same isolation batch was used for all experiments), in media supplemented with 50ng/mL M-CSF and various serum additives as follows: 10% heat-inactivated fetal bovine serum (hiFBS), 10% heat-inactivated human serum (hi huS), 10% non-hi huS, or 10% non-hi huS + 200μg/mL Compstatin. Human serum used for all conditions was from the same lot of pooled human AB serum (Fisher Scientific, Cat#: BP2525-100). For heat inactivation, serum was placed in a 56°C water bath for 30 min and stored in -20°C until use. Compstatin was purchased from Selleck chemical, Houston, TX, USA, Cat#: S8522. For ADAM17 inhibition, MΦ were treated with 10μg/mL PGN and concomitantly with 30μM of TAPI-0 or Marimastat (Tocris, McKinley Place, MN, USA, cat#’s: 5523, 2631).

### Preparation of Apoptotic Neutrophils

On the morning of Day 8, human neutrophils (PMN) were isolated by immunomagnetic negative selection from healthy donor buffy coats (EasySep Direct Human Neutrophil isolation kit, Cat#:19666). After isolation and counting, PMN were resuspended to 5×10^6^ cells/mL in IMDM without additives, labeled with eFluor670 according to manufacturer’s protocol (Thermo Fisher, Cat#: 64-0840-90), then exposed to 217.8 mJ/cm^2^ UV irradiation for 3 min to induce apoptosis. Irradiated PMN were then placed in a 37°C incubator with 5% CO_2_ for 4h. After 4hrs, an aliquot of PMN was stained with Annexin-V and PI (Annexin-V Apoptosis Detection kit, eBiosciences, Cat#:88-8005-74) and assessed for the frequency of Annexin V^+^/PI^−^ early apoptotic PMN by flow cytometry. All apoptotic cell preparations contained ≥70% early apoptotic cells.

### Efferocytosis Assay

On Day 8, labeled apoptotic PMN were co-incubated with MΦ to assess efferocytosis. Prior to adding apoptotic PMN, supernatants from cultured MΦ were replaced with 0.5 mL/well of IMDM without serum. The collected supernatant was stored at -80°C until use for detection of secreted cytokines. After 4hrs, irradiated e-fluor670-labeled PMN were added to UpCell plate-containing MΦ at a 5:1 apoptotic PMN:MΦ ratio and incubated for 1hr (2.5×10^6^/mL PMN:0.5×10^6^/mL MΦ). After 1hr, UpCell plates were gently washed two times using 1mL PBS per wash to remove unbound PMNs. UpCell plates were rested at room temperature for 10 min. to allow MΦ detachment. To ensure complete removal, wells were washed once with 1 mL ice cold PBS.

### Flow Cytometry

MΦ (0.5×10^6^/tube) were stained for flow cytometry in 5mL round-bottom polystyrene tubes (Corning, NY, USA, Cat#: 352058). After pelleting the cells at 1200rpm for 5 min, they were incubated for 10 min at room temperature with a combined Fc block solution (2μL Human Fc block [BD Biosciences, San Jose, CA, USA] + 3μL heat-inactivated human AB serum [Fisher Scientific, Waltham, MA, Cat#: BP2525-100] + 5μL Brilliant Stain buffer [Thermo Fisher, Waltham, MA, Cat#: 00-4409-42]) per tube. Primary antibodies were then added and after gentle mixing, incubated for 25 min on ice. After washing the cells with FACS Wash (2% FBS, 1mM EDTA in PBS), unconjugated antibodies were detected with secondary PE-conjugated antibody for 20 min on ice. After staining all samples were washed with 2mL FACS Wash, fixed in 1% paraformaldehyde in 0.15M NaCl, pH 7.4 and kept in 4°until analysis. Data was collected using an LSRII flow cytometer (BD Biosciences) and analyzed using FlowJo software (Treestar, Ashland, OR, USA, version 10.7.1). Number of events recorded were: 20,000/treatment for receptors and 40,000/treatment for efferocytosis assay. Antibodies and clones included in the efferocytosis cocktail were from BioLegend (San Diego, CA, USA): PE-conjugated anti-human CD11b (clone ICRF44), APC-Cy7-conjugated anti-human CD163 (clone GHI/61), PerCP-Cy5.5-conjugated anti-human CD206 (clone 15-2), AlexaFluor-488-conjugated anti-human CD64 (clone 10.1), or from BD Biosciences, BV421-conjugated anti-human CD66b (clone G10F5). For detecting surface receptor expression, efferocytic receptors were all PE-labeled and purchased from BioLegend, CD36 (clone 5-271), or purchased from R&D Systems as follows: DTK/TYRO3 (clone 96201), AXL (clone 108724), MERTK (clone 125518), αVβ3 (clone), or αVβ5 (clone P5H9), CD300f (clone UP-D2). Additional receptors were only available unconjugated and purchased from R&D Systems: anti-human CD300b (polyclonal goat IgG, Cat#: AF2879) and stained with secondary antibody (PE-conjugated anti-goat IgG, Cat#: F0107). Anti-human STABILIN-1 and anti-human STABILIN-2 primary antibodies (both polyclonal sheep IgG, Cat#: AF3825, Cat#: AF2645) were stained with a PE-conjugated secondary antibody (polyclonal anti-sheep IgG, Cat#: 5-001-A). Receptor expression was quantified using mean fluorescence intensity (MFI) for each respective treatment group.

### Measurement of soluble analytes

Supernatants from PGN-treated MΦ (24hrs in non-hi huS) were tested for soluble analytes (AXL, CD36, CD44, ICAM-1, LOX-1, MERTK, RAGE, TIM-3, TNF, and TYRO3) using a custom Procarta Luminex assay from Thermo Fisher (Waltham, MA, USA). Samples were diluted 1:5 or 1:10 in assay buffer and tested according to the manufacturer’s protocol. Results were read on a BioPlex 200 instrument and analyzed using BioPlex Manager software (BioRad, Hercules, CA, USA).

### Data analysis

All data are presented as mean ± SEM, with the number of independent donors indicated in the figure legends. Results were analyzed by: one-way or two-way ANOVA, or a mixed-effect analysis followed by a Dunnett’s or Sidak’s multiple-comparison post-hoc test (p < 0.05), a paired t-test, or a non-linear fit model as indicated in the figure legends. Graphs and statistics were generated in GraphPad Prism (GraphPad Prism, version 9.2.0, San Diego, CA, USA). * p<0.05, ** p<0.01, *** p<0.005, **** p<0.0001).

## RESULTS

### Human serum factors enhance human MΦ recognition of *Bacillus anthracis* PGN

To evaluate whether human serum opsonins are important for MΦ responses to peptidoglycan (PGN), we tested primary human MΦ for their capacity to produce cytokines in response to PGN in the presence of various serum conditions. We evaluated TNF and IL-10 production following exposure to 10μg/mL purified *B. anthracis* PGN in the presence of the following:10% heat-inactivated bovine serum (hi FBS), 10% heat-inactivated human serum (hi huS), 10% non-inactivated human serum (non-hi huS), and 10% non-inactivated human serum pre-treated with Compstatin, a specific inhibitor of C3b-mediated opsonization (non-hi huS+Comp). We observed negligible production of TNF and IL-10 in PGN-treated MΦ in the presence of hiFBS; however, in the presence of human serum, PGN induced large increases in both TNF and IL-10 concentrations in supernatants (**Figure 1A**).

### *Bacillus anthracis* PGN inhibits human MΦ efferocytosis of apoptotic PMN

Next, we tested whether exposure of M2-like MΦ to PGN impacts their capacity to efferocytose human apoptotic neutrophils, also termed polymorphonuclear leukocytes (PMN). MΦ were pre-treated with PGN for 24hrs in the presence of the various serum conditions shown in Figure 1A, then co-incubated with primary apoptotic PMNs for one hr, followed by assessment of efferocytosis using flow cytometry. Consistent with a human-serum opsonin requirement for recognition of PGN by human MΦ, efferocytosis was modestly impaired in the presence of hiFBS but profoundly impaired in the presence of all human serum conditions (**Figure 1B, C**). When directly compared, efferocytosis was slightly reduced in the presence of hi huS compared to non-hi huS (32 ± 4.9 SEM vs, 56.3 ± 5.2 SEM; p=0.0005), but inhibition of C3 showed no difference (non-hi huS+Comp, 55.6 ± 5.4 SEM vs, non-hi huS 56.3 ± 5.2 SEM p=0.99), indicating thermolabile opsonins, other than C3, aid PGN recognition (**Figure 1D**). We also performed a dose-response assay to find the 50% inhibitory concentration (IC_50_) of PGN on efferocytosis using MΦ from three representative donors **(Supplemental Figure 2)**. PGN elicited similar responses in both non-hi huS and non-hi huS + Compstatin in its capacity to inhibit efferocytosis (IC_50_ = 1.46μg/mL vs 1.49μg/mL), reinforcing that complement component C3 contributes little to PGN’s inhibition of efferocytosis in our system.

**Figure 2.**
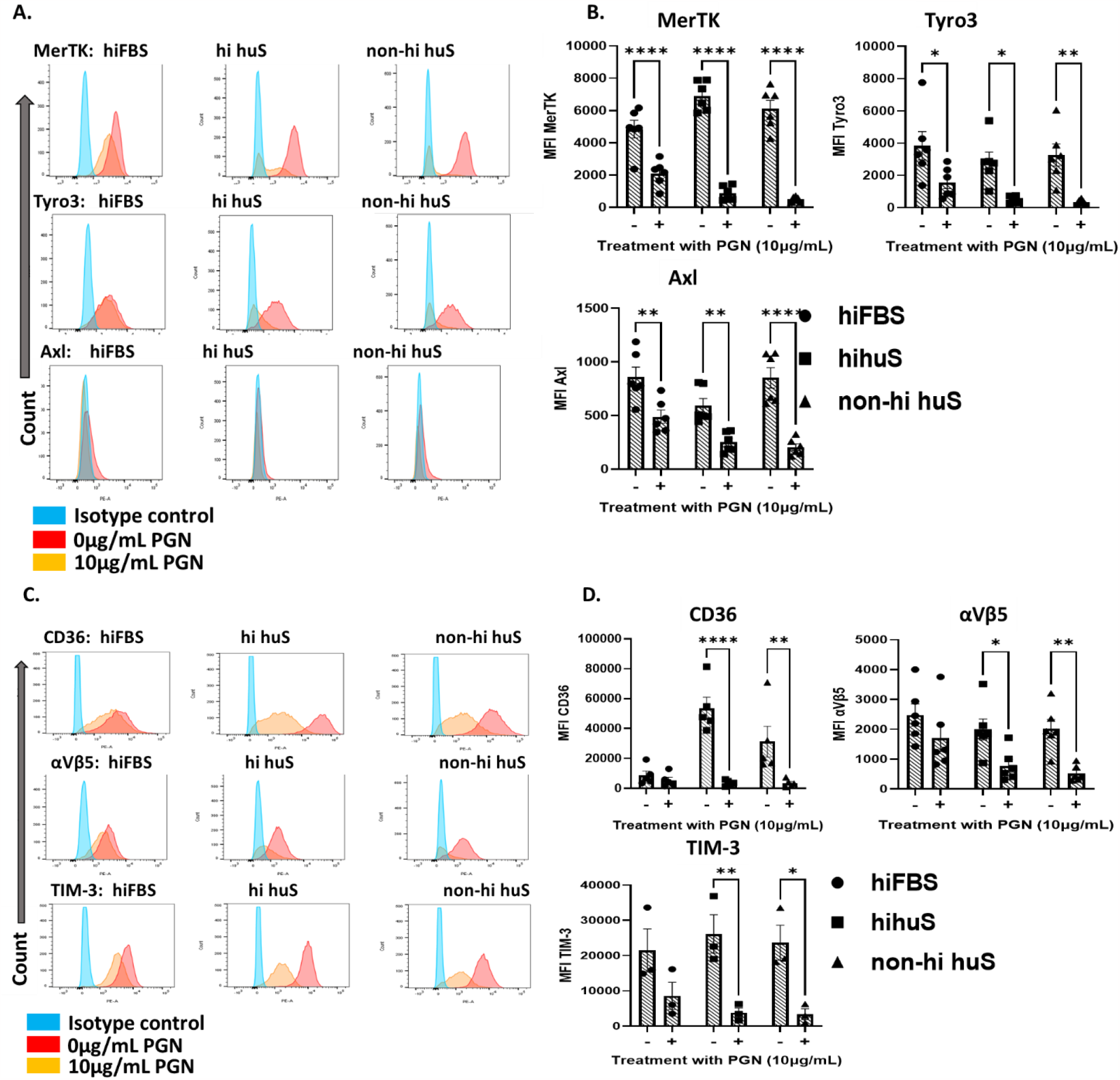
Human M2-like MΦ reduce efferocytic receptor expression in response to PGN. **(A, C)**. Representative histograms of receptor expression. **(B, D)** MFI of surface efferocytosis receptors from the indicated serum conditions from ≥3 independent donors analyzed by two-way ANOVA with Sidak’s multiple comparison.

### PGN treatment reduces cell surface expression of efferocytic receptors through proteolytic cleavage

Next, we investigated the cell surface expression of key efferocytosis signaling receptors in the presence of hiFBS, hi huS and non-hi huS. We measured a total of thirteen efferocytosis receptors, and found significant changes in six of them: TYRO3, AXL, MERTK, Integrin αVβ5, TIM-3, and CD36 (**Figure 2A-D**). The remaining seven receptors showed low expression levels or were unchanged following PGN treatment (**Supplemental Figure 3A-D**). All of the significantly reduced cell surface receptors, except integrin αVβ5, have been shown to be regulated via receptor cleavage^39^. Next, we measured levels of multiple efferocytosis-associated molecules using a custom bead-based multiplex assay (**Figure 3)**. A Disintegrin And Metalloproteases (ADAMs) are membrane-bound metalloproteases that act as sheddases to cleave various receptors from the cell membrane^40^. In particular, ADAM17 is one of the well-studied ADAM family members and is known to cleave a multitude of receptors including TNF26, TYRO3^41^, AXL^42^, MERTK^43^, CD36^44^ and TIM-3^45^, CD44^46^, ICAM-147, LOX-148, and RAGE49. Of the 10 ADAM17-mediated receptors/cytokines we measured, 6 increased in PGN-treated supernatants; however, soluble forms of TYRO3, AXL, MERTK, and TIM-3 were undetectable (**Figure 3A, B)**. We observed an inverse correlation between concentration of TNF and efferocytosis (**Figure 3C, D**); however, TNF blockade failed to prevent the PGN-induced defect (**Supplemental Figure 4**). Overall, PGN reduced the cell surface expression of six key efferocytosis receptors on human M2-like MΦ and increased multiple ADAM17 substrates, suggesting involvement of membrane-bound proteases.

**Figure 3:**
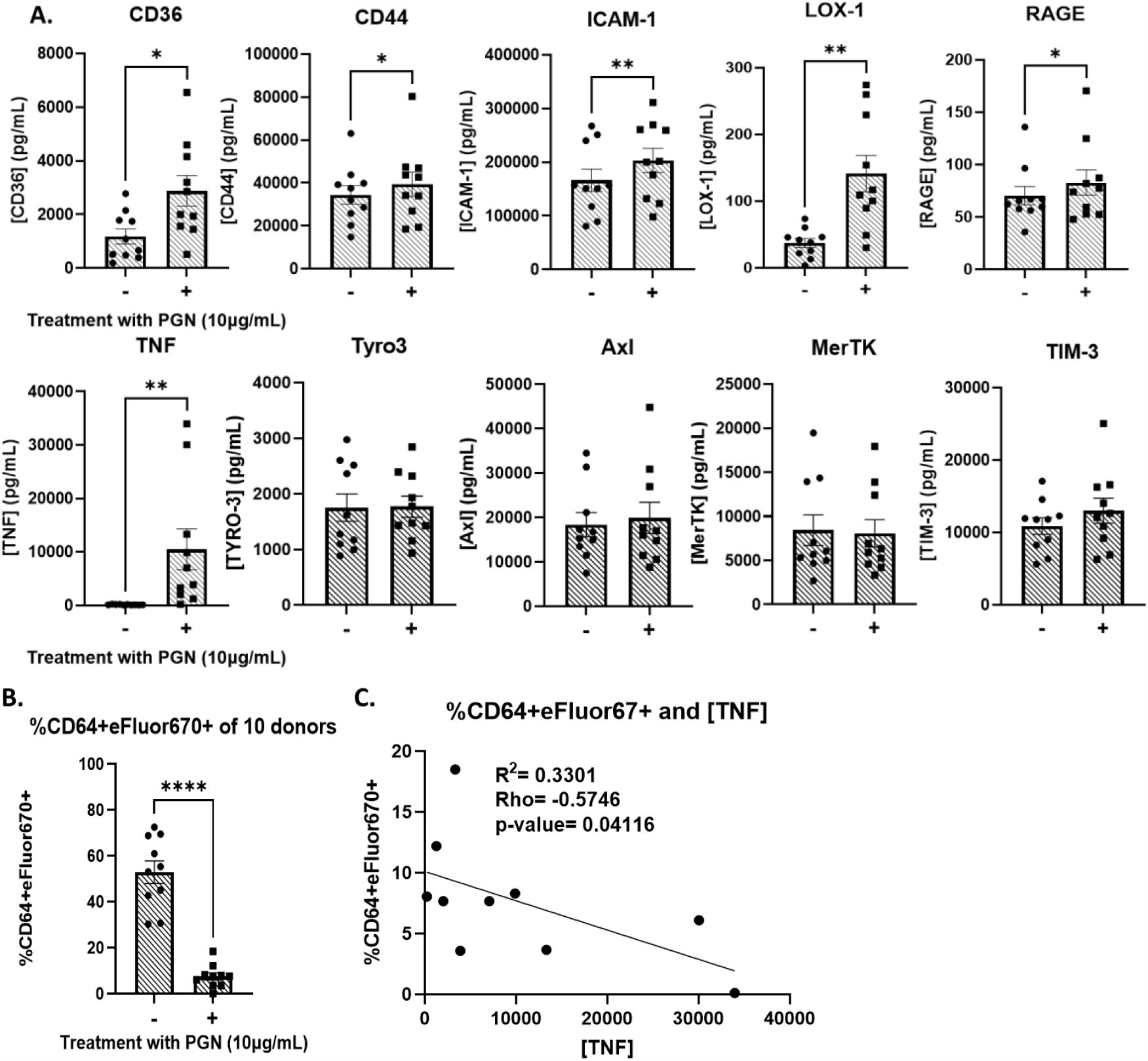
PGN-treated supernatants have increased levels of ADAM17 substrates. **A)** Concentrations of soluble receptors and cytokines from efferocytosis supernatants that are known to be cleaved by ADAM17. **B)** %CD64+eFluor670+ efferocytosis from 10 donors, PGN-treated values were used for the correlation analysis **C)** Pearson correlation and Linear Regression analysis between %efferocytosis and concentration of TNF. PGN values are from panel B. All analytes were measured from 24hr PGN-treated supernatants in the non-hi huS serum condition analyzed by paired t-tests.

**Figure 4:**
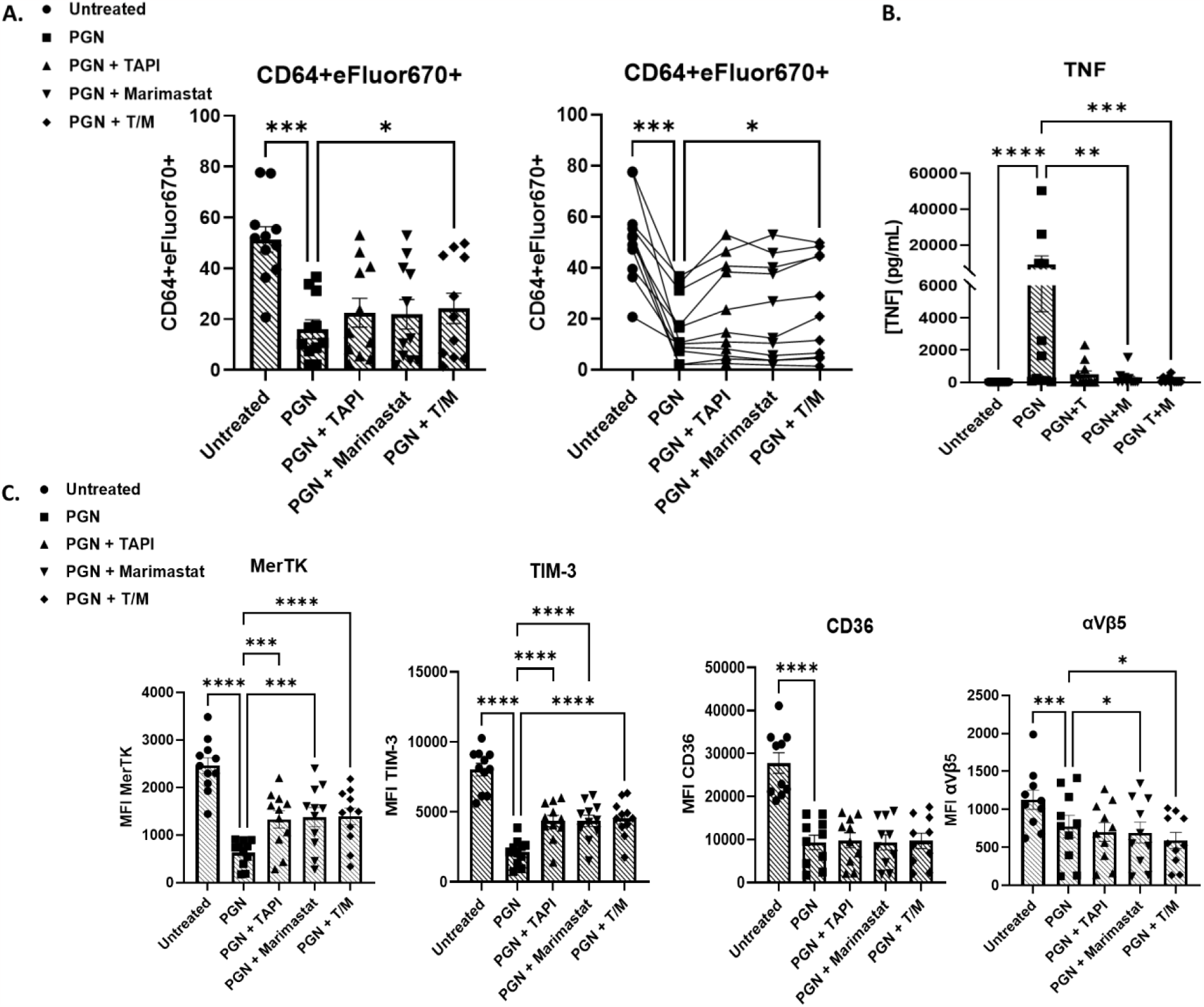
Pharmacological inhibition of ADAM17 partially restores surface receptor expression and efferocytic capacity. **A)** Mean % efferocytosis (left) or an alternative display of the data showing the same donor tracked across each treatment group (right). **B)** TNF concentrations from donor supernatants. **C)** Level of receptor expression on cell surface (MFI). Data are from 11 independent donors after 16hr pre-treatment with 10μg/mL PGN in the presence of 10% non-hi huS serum alone or in combination with ADAM17 inhibitors. A, B and C are paired data from the same donors

### Blockade of ADAM17 abolished TNF release but only modestly preserved efferocytic capacity and receptor expression

Regulation of ADAM17 activity is complex, based on its maturation and trafficking between the endoplasmic reticulum and Golgi, and its sub-cellular localization in lysosomal and recycling compartments^50, 51^. Using two small molecule inhibitors, TAPI-0 and Marimastat, we evaluated if blocking ADAM17 could restore efferocytosis and receptor expression. Though TAPI-0 and Marimastat follow a similar mechanism of inhibition, by chelating zinc ions in ADAM17’s active site, TAPI-0 is not membrane-permeable and inhibits the enzyme at the cell surface, while Marimastat can permeate cellular membranes^52^. First, we titrated TAPI-0 (1, 30, 60μM, IC_50_ = 0.1μM) and found no difference in efferocytosis among the three conditions (**Supplemental Figure 5**). Next, we tested either inhibitor alone, or in combination, and observed partial increases in MERTK and TIM-3 and a minor, though statistically significant increase in efferocytosis (**Figure 4A, C**). Following treatment with PGN, supernatants from cultures containing the ADAM17 inhibitor TAPI-0 alone, or in combination with Marimastat showed substantially reduced TNF levels indicating effective inhibition of enzyme activity (**Figure 4B)**.

## DISCUSSION

While *Bacillus anthracis* produces toxins that enable the pathogen to replicate to high numbers^53^, PGN is an important factor that contributes to late sepsis-like pathology^28, 29, 54^. During sepsis and other forms of inflammation, there is an accumulation of uncleared apoptotic cells and host DAMPs including nucleosomes^55, 56^, suggesting potential defects in efferocytosis. We previously showed that *B. anthracis* edema toxin can inhibit efferocytosis by human M2-like MΦ^27^. In the present study, we investigated the effect of PGN on human MΦ efferocytosis. Herein, we report that PGN decreases efferocytosis in an opsonin-dependent manner. We also provide evidence that MΦ recognition of PGN is independent of complement component C3, one of the most abundant and promiscuous serum opsonins, as TNF production was equivalent in the presence of hi huS compared to non-hi huS. In contrast, efferocytosis was less efficient in the presence of hi huS, and this could reflect denaturation of efferocytic bridge proteins present in human serum, C1q, or both, as efferocytosis was unaffected by inhibition of the complement component C3. These findings agree with previous reports showing that other human immune cells, generally require human serum opsonins for effective responses to polymeric PGN^57, 58^. However, human MΦ did not require complement component C3 to respond to PGN, which differs from human PMNs^37^.

To investigate potential mechanisms leading to defective efferocytosis, we measured cell-surface expression of key receptors that signal for apoptotic cell engulfment including: TAM (TYRO, AXL, MERTK), TIM (TIM-1, TIM-3), Integrin (αVβ3, αVβ5), CD300 (CD300b, CD300f), and Stabilin (STABILIN-1, STABILIN-2), as well as the thrombospondin receptor, CD36. Though the transmembrane receptor BAI-1 is considered to be an efferocytosis signaling receptor in mice^59^, a recent publication failed to find expression of this receptor at the protein or RNA level in human MΦ^60^; therefore, we excluded this receptor from our screening. In total, cell surface expression of six receptors were significantly reduced following exposure to *B. anthracis* PGN for 24hrs (TYRO3, AXL, MERTK, TIM-3, αVβ5, and CD36), suggesting that downregulation of these receptors might be responsible for PGN-mediated defects in our model.

One known mechanism of regulation, common to multiple efferocytosis receptors, is enzyme-mediated shedding. In humans, ADAM17 is known to cleave multiple efferocytosis receptors, resulting in their soluble forms^22^. Cleavage of receptors can prevent efferocytosis by reducing the cell-surface expression of the receptor, and soluble forms act as decoy receptors to bind phosphatidylserine sites and inhibit recognition of apoptotic cells by other MΦ. Previous work has shown lipopolysaccharide (LPS) inhibits efferocytosis in a TNF-dependent mechanism in unpolarized human MΦ^61^. Work by Michlewska et al noted IL-10 and TNF show opposing secretion profiles, interestingly our work demonstrated that both IL-10 and TNF increase following treatment with PGN. Our work revealed that TNF was not responsible for *B. anthracis* PGN-induced defective efferocytosis by human M2-like MΦ. Our data showed that both ADAM17 inhibition, which prevented TNF release, and TNF neutralization failed to substantially restore efferocytosis. Michalowski et al. pretreated non-polarized MΦ with LPS for an extended period of time (120hrs), resulting in much higher TNF levels; thus, differences between the studies may be attributable to differing MΦ activation states and differences in TNF levels. Consistent with PGN-induced sheddase activity, we observed significant increases in the soluble forms of CD36, as well as ICAM-1, CD44, LOX-1 and RAGE, key efferocytosis receptors that are cleaved by ADAM17.

To block ADAM17, we used two small-molecule inhibitors, TAPI-0 and Marimastat, and saw partial restoration of efferocytosis as well as a partial increase in MERTK and TIM-3 on the cell surface, however neither returned to baseline levels. Therefore, this study suggests the existence of additional mechanisms for downregulation of these receptors and for PGN-mediated impairment of efferocytosis, some of which may be transcriptional or post-translational. Interestingly, we did not observe changes in the levels of soluble TYRO3, AXL, MERTK or TIM-3, suggesting that *B. anthracis* PGN induces selective substrate cleavage and/or alternative regulation beyond cell-surface expression. Work by Maretzky et al have shown that ADAM17 can cleave different substrates when in the presence of rhomboid proteins 1 or 2 (iRhom1, iRhom2), demonstrating that when complexed together, iRhom proteins can modify ADAM17 substrate specificity^62, 63^. To the best of our knowledge substrate specificity as a mechanism of receptor regulation has yet to be shown for the TYRO, AXL, MERTK (TAM) family. Of note, the TAM family was demonstrated to be regulated through intracellular cleavage at their membrane-proximal region by gamma-secretase. Interestingly, the extracellular domain must be cleaved first, but Merilahti et al demonstrated that intracellular TAM family members could interact with cytosolic signaling intermediates and additionally, localize to the nucleus^64^. In addition, it has been noted in human MΦ from systemic lupus erythematosus (SLE) patients that the level of soluble receptor and membrane-bound forms of MERTK do not correlate^65^. This suggests nuanced regulation and partly explains our result of reduced cell-surface expression in PGN-treated MΦ yet an equal level of soluble receptor in PGN-and un-treated MΦ supernatant.

Although PGNs are present in both Gram+ and Gram^−^ bacteria, the PGN layer is much thicker in Gram+ species and therefore the pathological burden of PGN is expected to be higher during Gram+ infections such as anthrax. Furthermore, the structural heterogeneity among bacterial PGNs, such as variations in stem peptides and post-synthetic modifications of the glycan strands^66^, could lead to diverse biologically-active breakdown products that in turn alter downstream signaling and functional oucomes^32, 67, 68^. Further studies are required to determine whether other bacterial PGNs can inhibit the capacity of human MΦ to engulf apoptotic cells.

In summary, we report that PGN from *Bacillus anthracis* inhibits efferocytosis by human M2-like MΦ and that human serum is important for eliciting maximal response. We also demonstrate that PGN-mediated defects are partially due to ADAM17-mediated proteolysis of MERTK and TIM-3 receptors. Nevertheless, ADAM17 inhibition only partially restores PGN-induced efferocytic impairment, thus metalloprotease-independent mechanisms for regulating efferocytic receptors must also occur. Overall, these data show that bacterial factors can affect the capacity of MΦ to efferocytose and provide a new explanation for the accumulation of uncleared apoptotic cells in the context of late stage anthrax infection.

## ACKNOWLEDGEMENTS

The authors would like to thank Dr. Diana Hamilton, Jacob Bass and Dr. Linda Thompson at the OMRF Flow Cytometry Core for assistance with flow cytometry, and Dr. Susan Kovats for her helpful advice.

## AUTHOR CONTRIBUTIONS

JSM, ZP, and ADF conceived, designed, and supervised the experiments. JSM, ZP and JZ collected and analyzed the data. NP contributed to the reagents and materials (*B. anthracis* PGN isolation and validation of purity). JAJ and NR contributed reagents and materials (healthy donor recruitment and blood collection). CLD and CL isolated PMN and monitored apoptosis by flow cytometry. JSM, ZP and ADF wrote the manuscript. All authors revised the manuscript and approved the final version.

## SUPPLEMENTAL FIGURES

**Supplemental Figure 1:**
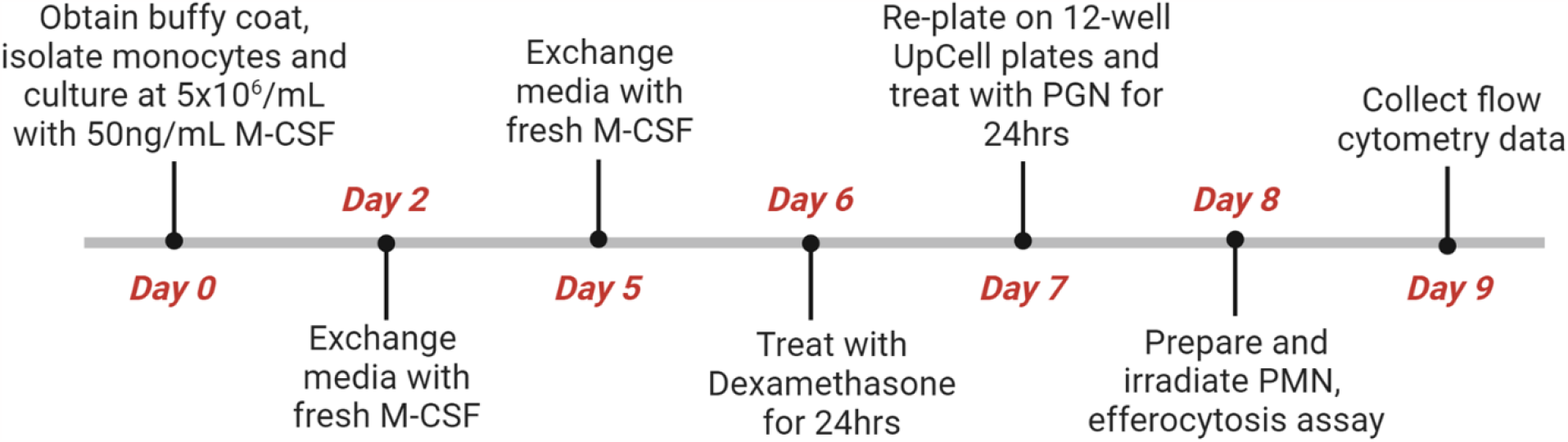
Overview of experimental setup. On Day 0 MΦ were isolated from buffy coat donors, adhered to plates for 1h, and cultured with M-CSF. On Day 2 and Day 5 half of the media was replaced with fresh complete media supplemented with 2x M-CSF to ensure differentiation. MΦ were polarized with Dexamethasone on Day 6, and 24hrs later, were re-plated onto 12-well UpCell plates and stimulated with various treatments. On Day 8 PMN were isolated, irradiated, co-cultured with MΦ for 1hr and subsequently stained for flow cytometry. PGN-treated supernatant was removed prior to PMN co-culture. On Day 9 samples were stained for flow cytometry, and data were collected (LSRII) and analyzed (FlowJo). Figure was created using Biorender, Template (9 Segments, Horizonal), by BioRender.com (2023). Retrieved from https://app.biorender.com/biorender-templates.

**Supplemental Figure 2:**
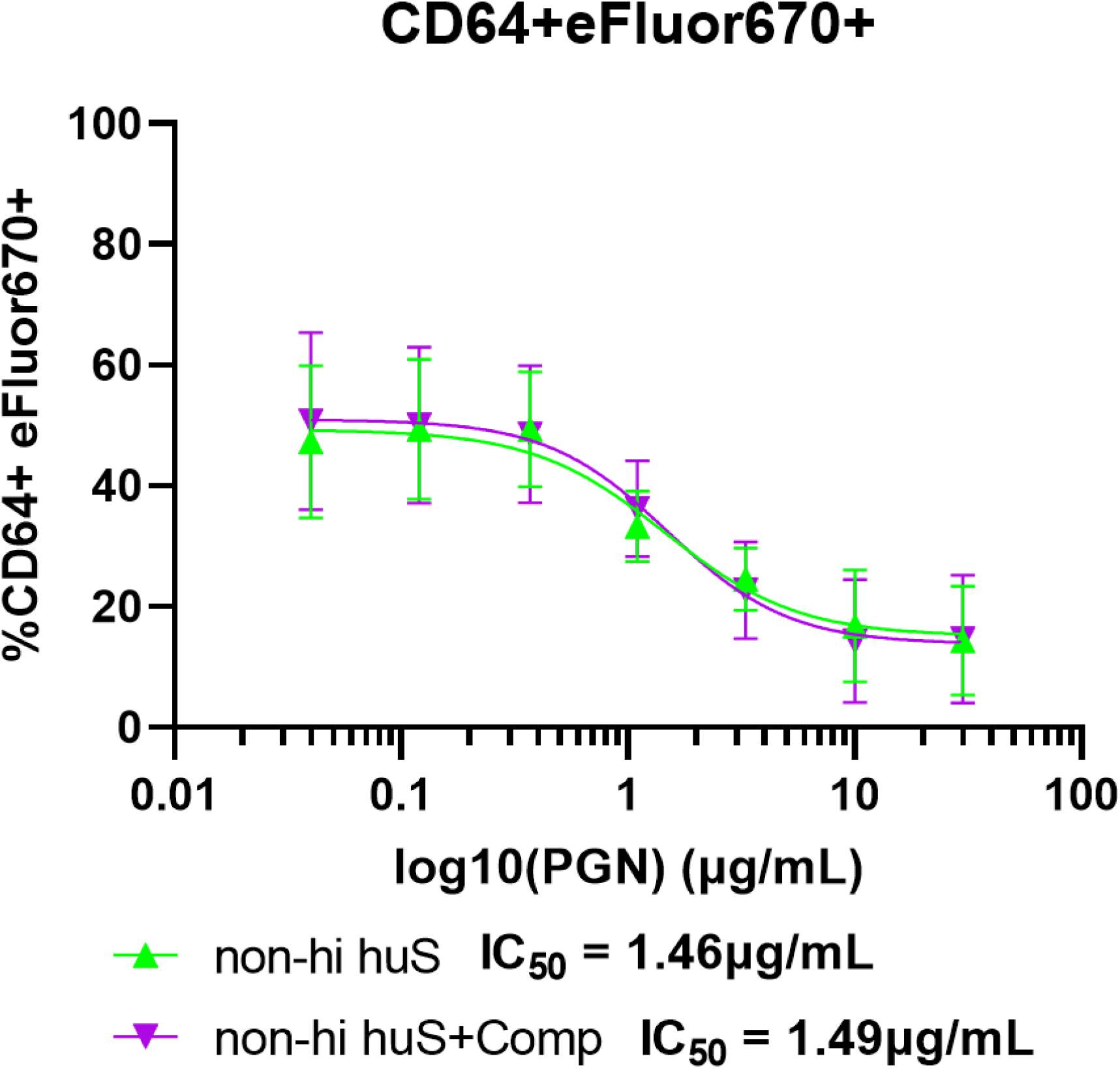
Dose response of PGN on MΦ efferocytosis from non-hi huS or non-hi huS+ Compstatin conditions from 3 independent donors (30μg/mL-0.04μg/mL). Data was log(10)-transformed and graphed using a non-linear fit model in Prism. Dose-response curve was used to calculate IC_50_.

**Supplemental Figure 3:**
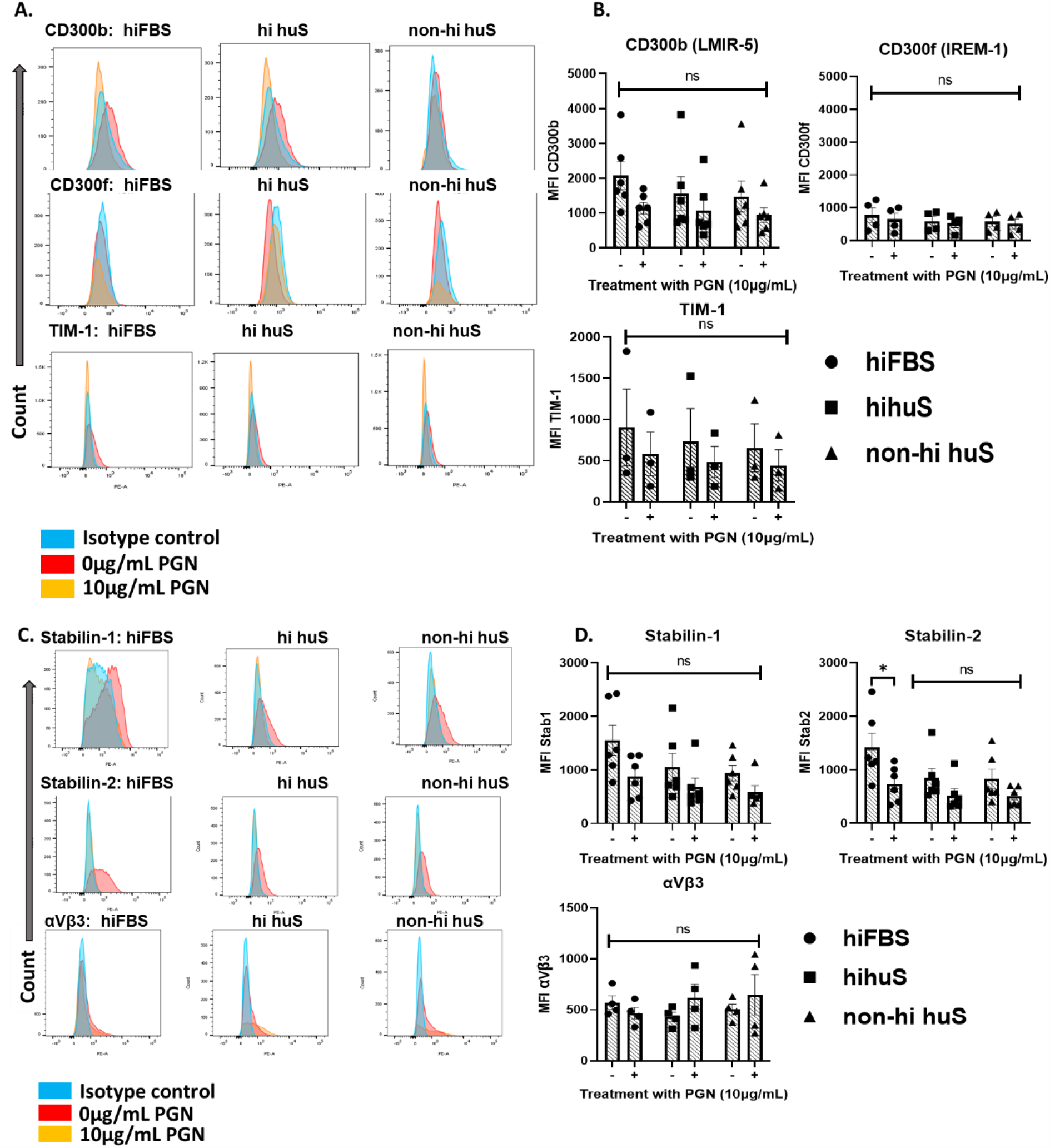
Efferocytosis receptors not affected by PGN treatment in non-hi huS in human M2-like macrophages. **(A, C)** Representative histograms of receptor expression. **(B, D)** MFI of surface efferocytosis receptors from the indicated serum conditions from ≥3 independent donors treated with 10μg/mL PGN for 24hrs and analyzed by two-way ANOVA with Sidak’s multiple comparison.

**Supplemental Figure 4:**
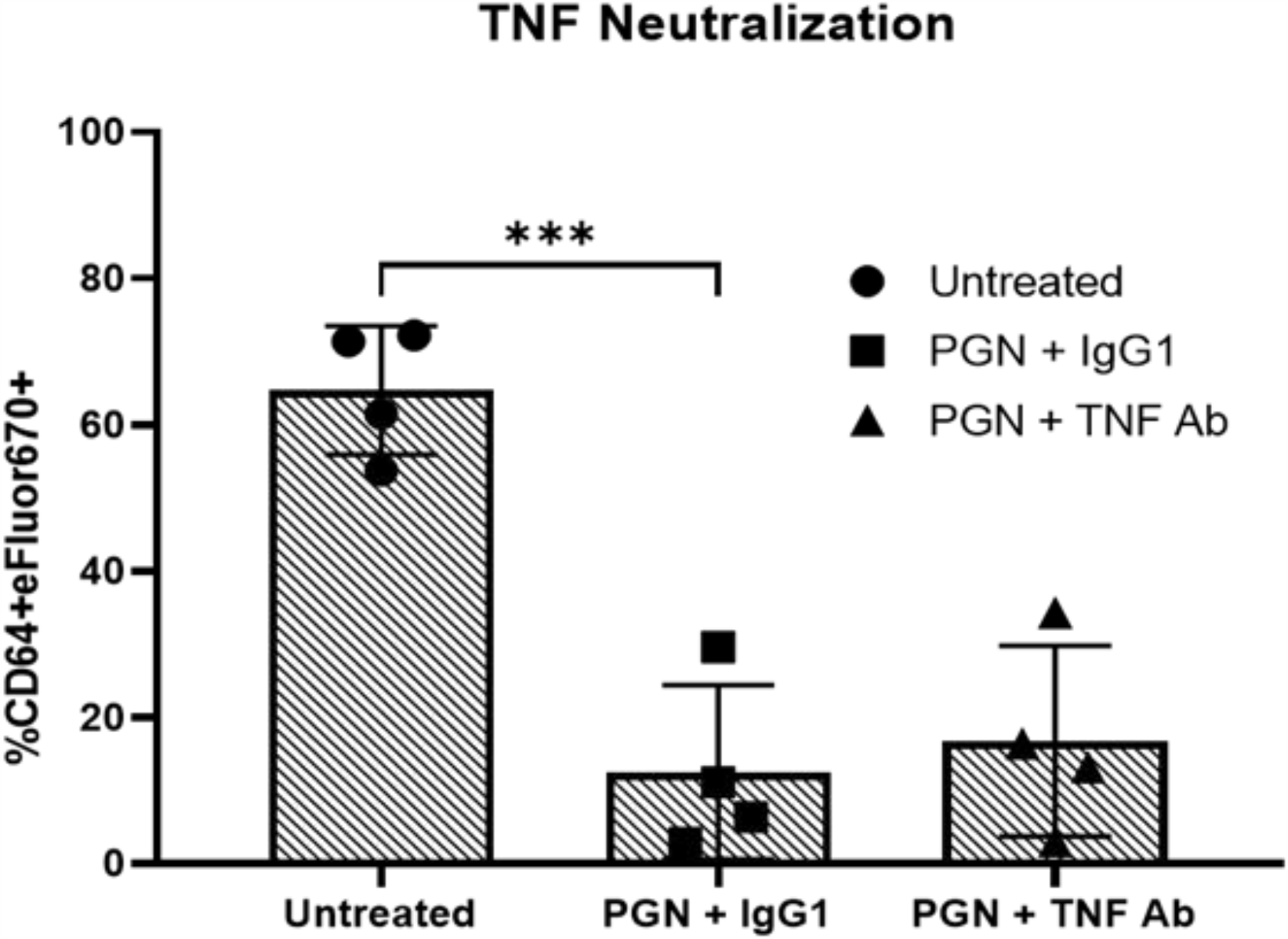
TNF neutralization does not rescue PGN-induced defects in efferocytosis. Percentage of CD64+ MΦ containing apoptotic eFluor670-labeled PMN treated with 6μg/mL of either an IgG isotype control (β-Galactosidase, InvivoGen Cat#: bgal-mab1) or human anti-TNF (IgG1, InvivoGen, Cat#: htnf-mab1) antibody in the presence of 10ug/mL PGN for 24 hrs. 4 independent donors analyzed by one-way ANOVA with Dunnet’s multiple comparison.

**Supplemental Figure 5:**
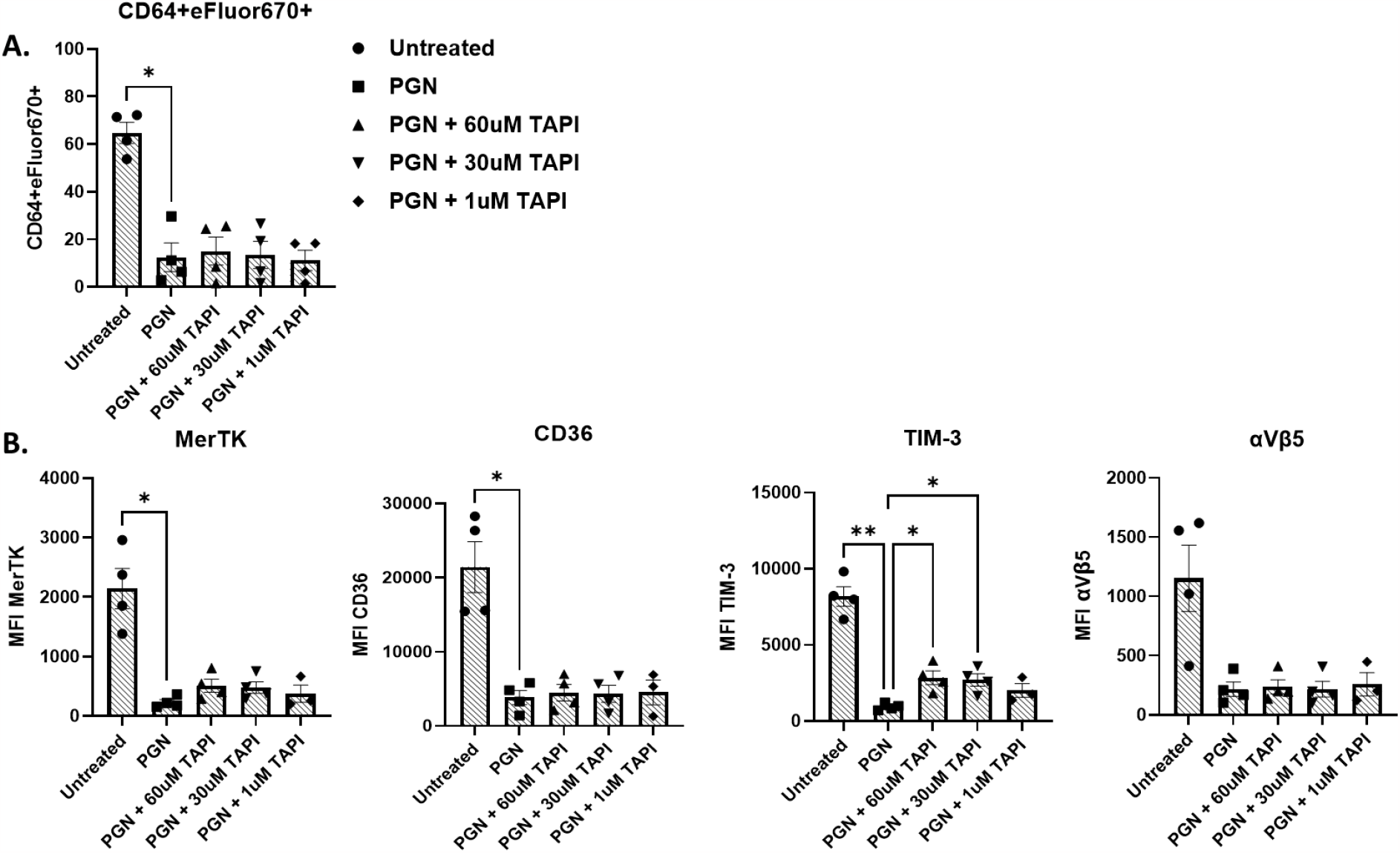
**A)** Percentage of CD64+ MΦ containing apoptotic eFluor670-labeled PMN from the non-hi huS condition in the presence of absence of 10ug/mL PGN and various doses of TAPI-0 (60, 30, 1μM). **B)** MFI of efferocytosis receptors from the same donors in Figure 5A. Data is from ≥3 independent and was analyzed using a mixed effects model with Sidak’s multiple comparison due to a missing data point for the 1μM TAPI condition.

## REFERENCES

1. Goel, A.K., Anthrax: A disease of biowarfare and public health importance. World J Clin Cases, 2015. 3(1): p. 20–33.

2. Rubinson, L.M.P., A. Corey, and D. Hanfling, Estimation of Time Period for Effective Human Inhalational Anthrax Treatment Including Antitoxin Therapy. PLoS Curr, 2017. 9.

3. Rudd, K.E., et al., Global, regional, and national sepsis incidence and mortality, 1990-2017: analysis for the Global Burden of Disease Study. Lancet, 2020. 395(10219): p. 200–211.

4. Cheng, Z., et al., The Critical Roles and Mechanisms of Immune Cell Death in Sepsis. Front Immunol, 2020. 11: p. 1918.

5. Li, Y., et al., Circulating Histones in Sepsis: Potential Outcome Predictors and Therapeutic Targets. Front Immunol, 2021. 12: p. 650184.

6. Xu, J., et al., Extracellular histones are major mediators of death in sepsis. Nat Med, 2009. 15(11): p. 1318–21.

7. Chen, Q., et al., Circulating nucleosomes as a predictor of sepsis and organ dysfunction in critically ill patients. Int J Infect Dis, 2012. 16(7): p. e558–64.

8. Zeerleder, S., et al., Elevated nucleosome levels in systemic inflammation and sepsis. Crit Care Med, 2003. 31(7): p. 1947–51.

9. Cummings, R.J., et al., Different tissue phagocytes sample apoptotic cells to direct distinct homeostasis programs. Nature, 2016. 539(7630): p. 565–569.

10. Baratin, M., et al., T Cell Zone Resident Macrophages Silently Dispose of Apoptotic Cells in the Lymph Node. Immunity, 2017. 47(2): p. 349–362 e5.

11. Gharib, S.A., et al., Transcriptional and functional diversity of human macrophage repolarization. J Allergy Clin Immunol, 2019. 143(4): p. 1536–1548.

12. Morioka, S., C. Maueroder, and K.S. Ravichandran, Living on the Edge: Efferocytosis at the Interface of Homeostasis and Pathology. Immunity, 2019. 50(5): p. 1149–1162.

13. Roberts, A.W., et al., Tissue-Resident Macrophages Are Locally Programmed for Silent Clearance of Apoptotic Cells. Immunity, 2017. 47(5): p. 913–927 e6.

14. Porcheray, F., et al., Macrophage activation switching: an asset for the resolution of inflammation. Clin Exp Immunol, 2005. 142(3): p. 481–9.

15. Dalli, J. and C.N. Serhan, Specific lipid mediator signatures of human phagocytes: microparticles stimulate macrophage efferocytosis and pro-resolving mediators. Blood, 2012. 120(15): p. e60–72.

16. Murray, P.J., et al., Macrophage activation and polarization: nomenclature and experimental guidelines. Immunity, 2014. 41(1): p. 14–20.

17. Roszer, T., Understanding the Mysterious M2 Macrophage through Activation Markers and Effector Mechanisms. Mediators of Inflammation, 2015. 2015.

18. Denning, N.L., et al., DAMPs and NETs in Sepsis. Front Immunol, 2019. 10: p. 2536.

19. Mehrotra, P. and K.S. Ravichandran, Drugging the efferocytosis process: concepts and opportunities. Nat Rev Drug Discov, 2022. 21(8): p. 601–620.

20. Greenlee-Wacker, M.C., Clearance of apoptotic neutrophils and resolution of inflammation. Immunol Rev, 2016. 273(1): p. 357–70.

21. Doran, A.C., A. Yurdagul, Jr., and I. Tabas, Efferocytosis in health and disease. Nat Rev Immunol, 2019.

22. Tajbakhsh, A., et al., Effect of soluble cleavage products of important receptors/ligands on efferocytosis: Their role in inflammatory, autoimmune and cardiovascular disease. Ageing Res Rev, 2019. 50: p. 43–57.

23. Tajbakhsh, A., et al., Potential diagnostic and prognostic of efferocytosis-related unwanted soluble receptors/ligands as new non-invasive biomarkers in disorders: a review. Mol Biol Rep, 2022. 49(6): p. 5133–5152.

24. Witas, R., et al., Defective Efferocytosis in a Murine Model of Sjogren’s Syndrome Is Mediated by Dysfunctional Mer Tyrosine Kinase Receptor. Int J Mol Sci, 2021. 22(18).

25. Cai, B., et al., MerTK cleavage limits proresolving mediator biosynthesis and exacerbates tissue inflammation. Proc Natl Acad Sci U S A, 2016. 113(23): p. 6526–31.

26. Bell, J.H., et al., Role of ADAM17 in the ectodomain shedding of TNF-alpha and its receptors by neutrophils and macrophages. J Leukoc Biol, 2007. 82(1): p. 173–6.

27. Pan, Z., et al., Bacillus anthracis Edema Toxin Inhibits Efferocytosis in Human Macrophages and Alters Efferocytic Receptor Signaling. Int J Mol Sci, 2019. 20(5).

28. Keshari, R.S., et al., Complement C5 inhibition protects against hemolytic anemia and acute kidney injury in anthrax peptidoglycan-induced sepsis in baboons. Proc Natl Acad Sci U S A, 2021. 118(37).

29. Popescu, N.I., et al., Peptidoglycan induces disseminated intravascular coagulation in baboons through activation of both coagulation pathways. Blood, 2018. 132(8): p. 849–860.

30. Takeuchi, O., K. Hoshino, and S. Akira, Cutting edge: TLR2-deficient and MyD88-deficient mice are highly susceptible to Staphylococcus aureus infection. J Immunol, 2000. 165(10): p. 5392–6.

31. Travassos, L.H., et al., Toll-like receptor 2-dependent bacterial sensing does not occur via peptidoglycan recognition. EMBO Rep, 2004. 5(10): p. 1000–6.

32. Girardin, S.E., et al., Peptidoglycan molecular requirements allowing detection by Nod1 and Nod2. J Biol Chem, 2003. 278(43): p. 41702–8.

33. Langer, M., et al., Neither Lys- and DAP-type peptidoglycans stimulate mouse or human innate immune cells via Toll-like receptor 2. PLoS One, 2018. 13(2): p. e0193207.

34. Stafford, C.A., et al., Phosphorylation of muramyl peptides by NAGK is required for NOD2 activation. Nature, 2022. 609(7927): p. 590–596.

35. Girton, A.W., et al., Serum Amyloid P and IgG Exhibit Differential Capabilities in the Activation of the Innate Immune System in Response to Bacillus anthracis Peptidoglycan. Infect Immun, 2018. 86(5).

36. Turner, S., et al., Gram-Positive Bacteria Cell Wall Peptidoglycan Polymers Activate Human Dendritic Cells to Produce IL-23 and IL-1beta and Promote T(H)17 Cell Differentiation. Microorganisms, 2023. 11(1).

37. Popescu, N.I., et al., C3 Opsonization of Anthrax Bacterium and Peptidoglycan Supports Recognition and Activation of Neutrophils. Microorganisms, 2020. 8(7).

38. Langer, M., et al., Neither Lys- and DAP-type peptidoglycans stimulate mouse or human innate immune cells via Toll-like receptor 2. Plos One, 2018. 13(2).

39. Moss, M.L. and D. Minond, Recent Advances in ADAM17 Research: A Promising Target for Cancer and Inflammation. Mediators Inflamm, 2017. 2017: p. 9673537.

40. Malemud, C.J., Inhibition of MMPs and ADAM/ADAMTS. Biochem Pharmacol, 2019. 165: p. 33–40.

41. Lee, I.J., et al., Monocyte and plasma expression of TAM ligand and receptor in renal failure: Links to unregulated immunity and chronic inflammation. Clin Immunol, 2015. 158(2): p. 231–41.

42. Orme, J.J., et al., Heightened cleavage of Axl receptor tyrosine kinase by ADAM metalloproteases may contribute to disease pathogenesis in SLE. Clin Immunol, 2016. 169: p. 58–68.

43. Thorp, E., et al., Shedding of the Mer tyrosine kinase receptor is mediated by ADAM17 protein through a pathway involving reactive oxygen species, protein kinase Cdelta, and p38 mitogen-activated protein kinase (MAPK). J Biol Chem, 2011. 286(38): p. 33335–44.

44. Driscoll, W.S., et al., Macrophage ADAM17 deficiency augments CD36-dependent apoptotic cell uptake and the linked anti-inflammatory phenotype. Circ Res, 2013. 113(1): p. 52–61.

45. Moller-Hackbarth, K., et al., A disintegrin and metalloprotease (ADAM) 10 and ADAM17 are major sheddases of T cell immunoglobulin and mucin domain 3 (Tim-3). J Biol Chem, 2013. 288(48): p. 34529–44.

46. Hart, S.P., et al., CD44 regulates phagocytosis of apoptotic neutrophil granulocytes, but not apoptotic lymphocytes, by human macrophages. J Immunol, 1997. 159(2): p. 919–25.

47. Wiesolek, H.L., et al., Intercellular Adhesion Molecule 1 Functions as an Efferocytosis Receptor in Inflammatory Macrophages. Am J Pathol, 2020. 190(4): p. 874–885.

48. Murphy, J.E., et al., LOX-1 scavenger receptor mediates calcium-dependent recognition of phosphatidylserine and apoptotic cells. Biochem J, 2006. 393(Pt 1): p. 107–15.

49. Friggeri, A., et al., Participation of the receptor for advanced glycation end products in efferocytosis. J Immunol, 2011. 186(11): p. 6191–8.

50. Soond, S.M., et al., ERK-mediated phosphorylation of Thr735 in TNFalpha-converting enzyme and its potential role in TACE protein trafficking. J Cell Sci, 2005. 118(Pt 11): p. 2371–80.

51. Dusterhoft, S., et al., Status update on iRhom and ADAM17: It’s still complicated. Biochim Biophys Acta Mol Cell Res, 2019. 1866(10): p. 1567–1583.

52. Lorenzen, I., et al., Control of ADAM17 activity by regulation of its cellular localisation. Sci Rep, 2016. 6: p. 35067.

53. Hutt, J.A., et al., Lethal factor, but not edema factor, is required to cause fatal anthrax in cynomolgus macaques after pulmonary spore challenge. Am J Pathol, 2014. 184(12): p. 3205–16.

54. Qiu, P., et al., Bacillus anthracis cell wall peptidoglycan but not lethal or edema toxins produces changes consistent with disseminated intravascular coagulation in a rat model. J Infect Dis, 2013. 208(6): p. 978–89.

55. Sharma, S.K. and G. Naidu, The role of danger-associated molecular patterns (DAMPs) in trauma and infections. J Thorac Dis, 2016. 8(7): p. 1406–9.

56. Elliott, M.R., K.M. Koster, and P.S. Murphy, Efferocytosis Signaling in the Regulation of Macrophage Inflammatory Responses. J Immunol, 2017. 198(4): p. 1387–1394.

57. Sun, D., et al., Anti-peptidoglycan antibodies and Fcgamma receptors are the key mediators of inflammation in Gram-positive sepsis. J Immunol, 2012. 189(5): p. 2423–31.

58. Iyer, J.K., et al., Inflammatory cytokine response to Bacillus anthracis peptidoglycan requires phagocytosis and lysosomal trafficking. Infect Immun, 2010. 78(6): p. 2418–28.

59. Park, D., et al., BAI1 is an engulfment receptor for apoptotic cells upstream of the ELMO/Dock180/Rac module. Nature, 2007. 450(7168): p. 430–U10.

60. Hsiao, C.-C., et al., Macrophages Do Not Express the Phagocytic Receptor BAI1/ADGRB1. Frontiers in Immunology, 2019. 10(962).

61. Michlewska, S., et al., Macrophage phagocytosis of apoptotic neutrophils is critically regulated by the opposing actions of pro-inflammatory and anti-inflammatory agents: key role for TNF-alpha. Faseb Journal, 2009. 23(3): p. 844–854.

62. Maretzky, T., et al., iRhom2 controls the substrate selectivity of stimulated ADAM17-dependent ectodomain shedding. Proc Natl Acad Sci U S A, 2013. 110(28): p. 11433–8.

63. Zhao, Y., et al., Identification of Molecular Determinants in iRhoms1 and 2 That Contribute to the Substrate Selectivity of Stimulated ADAM17. Int J Mol Sci, 2022. 23(21).

64. Merilahti, J.A.M., et al., Genome-wide screen of gamma-secretase-mediated intramembrane cleavage of receptor tyrosine kinases. Mol Biol Cell, 2017. 28(22): p. 3123–3131.

65. Zhu, H., et al., The expression and clinical significance of different forms of Mer receptor tyrosine kinase in systemic lupus erythematosus. J Immunol Res, 2014. 2014: p. 431896.

66. Vollmer, W., D. Blanot, and M.A. de Pedro, Peptidoglycan structure and architecture. FEMS Microbiol Rev, 2008. 32(2): p. 149–67.

67. Wolfert, M.A., A. Roychowdhury, and G.J. Boons, Modification of the structure of peptidoglycan is a strategy to avoid detection by nucleotide-binding oligomerization domain protein 1. Infection and Immunity, 2007. 75(2): p. 706–713.

68. Bersch, K.L., et al., Bacterial Peptidoglycan Fragments Differentially Regulate Innate Immune Signaling. ACS Cent Sci, 2021. 7(4): p. 688–696.

